# Forest vertical and horizontal temperature similarity drives arthropod communities in a managed temperate forest

**DOI:** 10.1101/2025.09.03.674010

**Authors:** Orsi Decker, Kerstin Pierick, Dominik Seidel, Christian Ammer, Bernhard Schuldt, Jörg Müller

## Abstract

Manipulating the canopy structure is the core tool of silviculture operation, and with that, changing the light availability alters temperature dynamics from the forest floor to the canopy. This should affect communities of ectothermic organisms such as insects, but we lack information on insect distributions in the complex 3D space of forests. Therefore, we set up temperature loggers and insect traps vertically (flight-interception traps) and horizontally (pitfall traps) in forests with experimental thinning and gap felling 8 years after the intervention. By metabarcoding, we identified ∼10,600 Operational Taxonomic Units (OTUs) from 44 orders including ∼2450 arthropods assigned to species in our 426 samples. Arthropod community similarity matrices were quantified along the Hill numbers accounting for rare to dominant species and under consideration of incomplete samples.

Arthropod communities were shaped by stratification (height above ground 0 m, 2 m, 10 m, 15 m), and by temperature similarity. Average nighttime temperature was the most important temperature variable for overall arthropod community similarity metrics. Restricted to flight interception traps, flying insect communities responded to daily temperature maximum and nighttime average temperature. Restricted to pitfall traps, on the other hand, arthropod communities were shaped by the overall temperature metric only when focusing on rare species. Additionally, all communities were strongly affected by season. Our results implies that management interventions establish different temperature heterogeneity within forest patches, which ultimately could drive species community similarity when including all arthropods in the area between forest floor and canopy.

## Introduction

Forest management is undoubtedly changing forest structure and biodiversity (Paillet et al. 2010). Management most often manipulates canopy opening and with that, light availability, which alters the temperature under canopy (Ehbrecht et al. 2019, Horváth et al. 2023, Menge et al. 2023, Brůna et al. 2024). Forest canopies are regarded as important buffers to climate change induced warming (De Frenne et al. 2019, De Frenne et al. 2021), however, measurements under the canopy are still scarce (but see del Pliego et al. 2016, Kovács et al. 2024, Pierick et al., 2025). Management not only changes the canopy, but ground structural complexity via altering the herb layer, litter and deadwood amount (Köhl et al. 2008, Paletto et al. 2014, Wang et al. 2021). Given that arthropods, the most diverse animal group in forests, are ectothermic, therefore likely to respond to light availability and temperature fluctuations not only on species-level (Lindman et al. 2022, 2023), but on a community-level (Sallé et al. 2021). Arthropods are widely using almost all microhabitats found under the canopy; thus, it is timely to investigate how canopy manipulations alter temperature dynamics and consequently shape arthropod communities inhabiting the forest floor and the space between the ground and canopy.

Microclimate is increasingly regarded as an important variable both in ecology and conservation (Hof et al. 2011, Lembrechts et al. 2019, Kemppinen et al. 2024, Kerr et al. 2025), given that species experience environmental variables in a resolution similar to their size (Wild et al. 2019, Pincebourde et al. 2021). However, the relationship and dynamics between several microclimate within a given area, such as a forest stand is still not well studied (but see Pierick et al., 2025). While some management interventions are successfully used to increase local biodiversity (Sebek et al. 2013, Doerfler et al. 2018, Thorn et al. 2020), the community responses to the possible microclimate changes following an intervention are often neglected. Management interventions could cause dramatic changes in the structure and microclimate of microhabitats arthropods use, such as deadwood addition or removal, damage to the shrub and herb layer, and canopy opening. As ectotherm organisms, with limited thermoregulation (Huey and Kingsolver 1989), arthropods might respond immediately and dramatically to temperature changes (Roitberg and Mangel 2016) induced by management. Forests in general offer a large spectrum of microclimates both spatially (Sears et al. 2011) and temporally (Zellweger et al. 2019), therefore arthropods are likely to have various available habitats within their thermal niche (Heidrich et al. 2020). However, forest management changes the thermal properties of a forest, especially under the canopy (Horváth et al. 2023), which ultimately modifies the available thermal niches for whole arthropod communities.

Vertical and horizontal temperature dynamics between the ground and the top of the canopy are most often overlooked and especially data on the vertical layers of a forest is scarce (Lenoir et al. 2017, De Frenne et al. 2021). Layers from the ground to the canopy offers a highly variable temperature pattern at various scales, such as created by the exposure of stems and leaves to UV radiation (Holden et al. 2011, Sears et al. 2011), which is likely to be important for animals foraging in this stratum (Ruchin 2023). Arthropods using the canopy are highly diverse given the resources canopy biomass offers (Müller et al. 2018, Procházka et al. 2018) and the largely enemy-frees space in this habitat (Wardhaugh 2014). However, sampling species in the different vertical layers is a challenge, especially in forest gaps, therefore mostly done with methods that pool arthropods from all layers (such as fogging). Studies from mostly tropical systems show that arthropods in the canopy are often heavily stratified depending on their foraging behaviours, life stages and the seasonality of breeding (Wardhaugh et al. 2014, Nakamura et al. 2017, Weiss et al. 2019, Urban-Mead et al. 2021). Therefore, management is likely to change the conditions for many species and whole communities, but rarely studied in temperate forests. Similarly, the ground surface in a forest is consisting of many living, abiotic and decomposing structures, such as shrubs and regenerating vegetation, large boulders or deadwood, which all have their individual temperature dynamics (Sears et al. 2011, Pincebourde and Woods 2020). Ground-living arthropods therefore are likely to use most of the existing forest floor structures depending on their size and thermal optimum, and the dynamics between these microclimates cannot be ignored.

Describing microclimates on a fine scale became possible in the past decade with fast-evolving devices, which log temperatures continuously (such as Wild et al. 2019) and models to process such data (reviewed in Kemppinen et al. 2024). In most descriptive studies, microclimate is considered on a low resolution (one logger per study site), which often fails to explain the temperature dynamics between forest structures (De Frenne et al. 2021, Kerr et al. 2025). This leads to a mismatch between the temperature of the study site (low-resolution) and the temperatures organisms experience (high-resolution). Nevertheless, studies show that diversity increases with the increasing available microclimates (Heidrich et al. 2020, Graf et al. 2022, Lettenmaier et al. 2022), however studies most often cover only arthropods inhabiting the forest floor, but arthropods found between the ground and the canopy are rarely studied in relation to microclimate(Nakamura et al. 2017). To address the lack of knowledge linking fine scale temperature metrics and arthropod community composition including communities from the forest floor to the canopy, we investigated the relationship between temperature and whole-community similarity in arthropods in a vertical and horizontal space. Following Pierick et al. (2025), we expect forest gaps to offer a higher microclimate variability compared to the thinning treatment, where canopy is still closed; and communities to be more similar to each other within similar temperature conditions.

## Materials and methods

### Study sites

The study was conducted in the managed areas of the Bavarian National Park, Germany on a small subset of plots which were established within the project BETA-FOR (Fig. 1a). BETA-FOR is a field experiment investigating the impact of forest structure manipulations on biodiversity and ecosystem functioning (Müller et al. 2023).). In these plots, 30% of the stand basal area was removed from treated plots, either aggregated in the center of the plot (“Gap”) or evenly distributed throughout the plot (“Thinning”), similar to forest management operations (Fig. 1b). The thinning and canopy openings were established in 2016 and the resulting opening in the Gap treatments had diameters of approximately 30 m. Both Gap and Thinning had two sub-treatments: entire trees were removed or standing deadwood of 5 m retained. For the current study, 2 geographical locations were used with two replicates of each treatment, resulting in 8 study plots. The plots are in temperate mountainous areas (average 931 m a.s.l.) where European beech (*Fagus sylvatica*) is the dominant species, reflecting a Central-European managed forest system. The full study including temperature dynamics from all locations can be found in Pierick et al., 2025.

**Figure 1.**
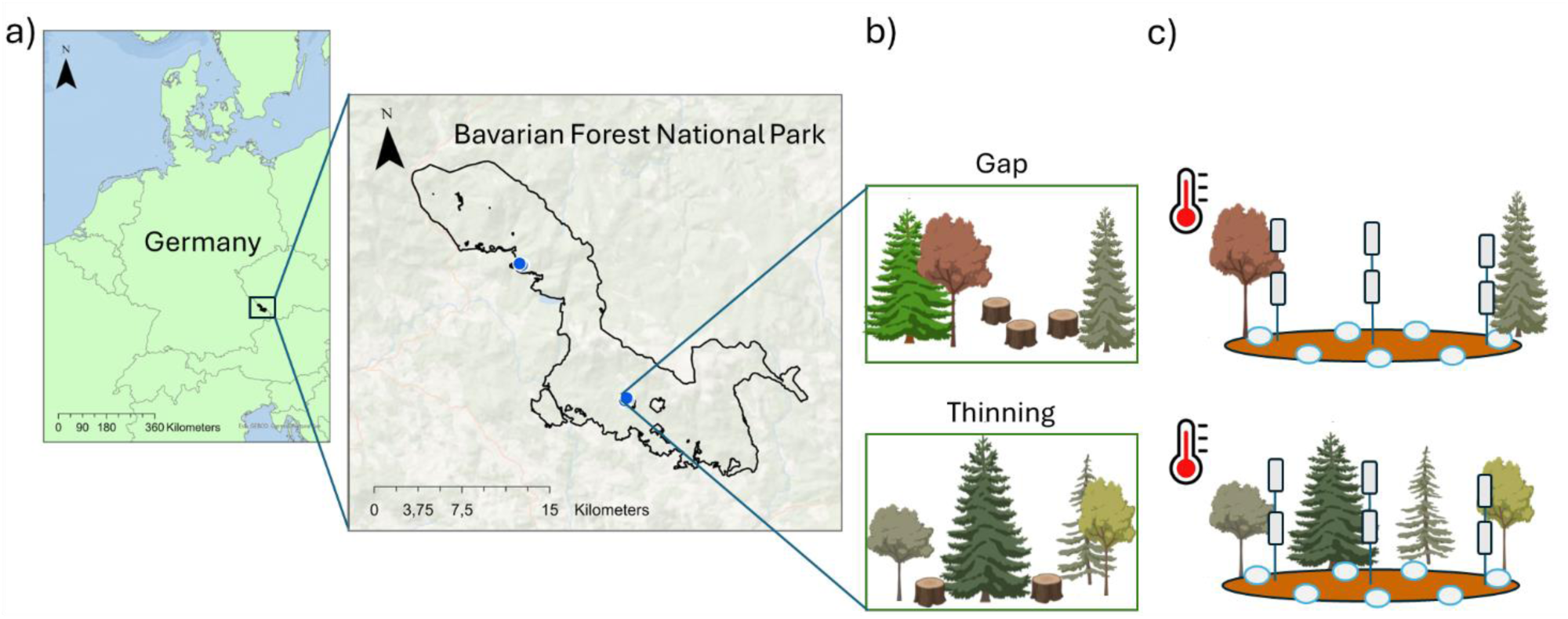
a) The location of study sites in the Bavarian National Park, Germany, where b) two main management types, Gap and Thinning were monitored for ground and canopy temperature and arthropods using c) flight-interception (rectangles) and pitfall traps (round dishes).

### Vertical temperature measurements

For measuring air temperatures under the canopy, three vertical ropes were installed (either directly fixed to branches or to horizontal ropes) in the canopy of the plots (Fig. 1c). One vertical rope was always positioned directly at the plot center, the other two 7.5 m north and south from the central rope, respectively. Each rope held three loggers, summing up to nine loggers per plot and a total of 72 devices in this study. The lowest logger was installed at 2 m, in the center of the plot, the uppermost as high as the conditions allowed, and the middle logger with approximately equal distances between the upper and lower ones, but generally aiming loggers at 2, 5, 10, 15 meters high. The maximum sensor heights per plot ranged from 12.5 m to 24.2 m. Onset Hobo MX2301A was used for vertical temperature monitoring (Onset, Bourne, USA) with TX Cover sun shields (Technoline, Bernburg, Germany) to protect them from direct solar radiation. We measured temperature in 30-minute intervals.

### Horizontal temperature measurements

To measure temperatures near the forest floor, 19 standard Tomst TMS-4 devices (Wild et al. 2019) were installed per plot, 152 loggers in total. The loggers were arranged in a hexagonal grid around the plot center, always 6 m from each other, gaining a maximal distance of 24 m for loggers at opposed corners of the grid (Fig. 1c). TMS-4 loggers measure temperatures 6 cm beneath the soil surface, near the soil surface (∼2 cm above), and 15 cm above the soil surface (Wild et al., 2019), but for the current study, we only used temperatures measured at 15 cm above soil surface. The measurement interval was set to 15 minutes.

### Arthropod sampling

Arthropods were sampled next to a subset of temperature loggers. Three flight-interception traps were installed under the two highest temperature loggers each at ∼10 m and ∼15 m height, and one flight-interception trap was installed at 2 m height at the center of the plot. This resulted in six flight-interception traps per study plot. Additionally, seven pitfall traps were installed next to Tomst loggers on the ground to represent the microhabitats of a given plot and aiming to install pitfall traps under the flight-interception traps. For example, most pitfall traps were under the canopy opening, but also in a position where the canopy was closed above the pitfall traps.

All arthropod traps were emptied every 14 days between mid-June and the end of August, 2024; however, 20 flight-interception traps could not be emptied on all occasions correctly either because of ropes being stuck or because of branches, and ropes collided following storms. As a result, the study includes 56 flight-interception traps and pitfall traps each with 4 time periods, equal to 428 arthropod samples (with 20 samples discarded or missing).

### Arthropod DNA metabarcoding

Arthropod samples were fractionated according to their body size into two categories: small and large species (or body parts) by sieving them through an 8mm sieve. This step was done to control for differences in biomass and to increase the identification of rare and small species (Morinière et al. 2016, Elbrecht et al. 2017). Following the fractionating step, the preservative ethanol was removed, and the mixed arthropod samples were dried overnight in a 60–70°C oven to eliminate residual ethanol. The dried arthropods were then homogenized using stainless steel beads in a FastPrep 96 system (MP Biomedicals). DNA extraction from all samples was conducted by incubating them in a 90:10 solution of animal lysis buffer (buffer ATL, Qiagen DNEasy tissue kit, Qiagen, Hilden, Germany) with 10% proteinase K following the standard Qiagen DNeasy protocol. Following an overnight incubation at 56°C, samples were cooled to room temperature. DNA was subsequently extracted from 200 μL aliquots using the DNEasy blood & tissue kit (Qiagen) according to the manufacturer’s instructions. Amplicon PCRs employed 5 μL of extracted genomic DNA, Plant MyTAQ (Bioline, Luckenwalde, Germany), and HTS-adapted mini-barcode primers targeting the mitochondrial Cytochrome-Oxidase subunit I (CO1-5P) region (mlCOIntF/ dgHCO2198) (Morinière et al. 2016, Leray and Knowlton 2017, Hausmann et al. 2020, Uhler et al. 2021). Amplification success and fragment length (313 bp) were confirmed via gel electrophoresis. Cleaned amplicons were resuspended in 50 μL molecular water. Illumina Nextera XT indices (Illumina Inc., San Diego, USA) were added in a secondary PCR using the same annealing temperature but limited to seven cycles. Ligation success was verified through gel electrophoresis, and DNA concentration measured with a Qubit fluorometer (Life Technologies, Carlsbad, USA). Samples were then pooled into 40 μL equimolar pools at 100 ng each, purified using MagSi-NGSprep Plus beads (Steinbrenner Laborsysteme GmbH, Wiesenbach, Germany), and eluted in a final volume of 20 μL. HTS was performed on mutiple Illumina NextSeq 2000 (v3 chemistry, 2*300 bp, 600 cycles) runs with a target of 250,000 paired-end reads per sample. Six controls were included on each 96-well plate: 2 DNA-extraction control, 2 PCR control and 2 ligation control to account for contamination or false positive results. To address uneven sample coverage, we applied a normalization approach that ensured each sample was analysed in proportion to its retained sequencing output, avoiding artificial rarefaction. This method preserves as much biological information as possible while maintaining comparability across samples. Importantly, all samples were processed at full sequencing depth rather than being subsampled to a uniform coverage threshold.

### Bioinformatics

Paired-end reads were merged using the USEARCH suite’s-fastq_mergepairs utility (v11.0.667_i86linux32) (Edgar 2010) with the following parameters:-fastq_maxdiffs 99, - fastq_pctid 75, and-fastq_trunctail 0. Adapter sequences were trimmed using CUTADAPT v5.0 (Martin 2011), with untrimmed sequences removed via the--discard-untrimmed option. Subsequent pre-processing—including quality filtering, dereplication, chimera removal, and clustering—was carried out with VSEARCH v2.30.1 (Linux x86 64-bit) (Rognes et al. 2016). Initial quality filtering employed--fastq_filter with parameters--fastq_maxee 1 and –minlen 300. The minimum length cutoff was chosen based on the expected amplicon size (313 bp) and downstream quality control considerations. After filtering, each sample retained ∼40,000 reads, with ∼30,000 reads remaining after dereplication for further analysis. Dereplication was performed using--derep_fulllength with the options--sizeout and--relabel Uniq, first at the sample level and then pooled across samples, with singletons excluded as noise. Reads were pre-clustered at 98% identity using the centroids algorithm, after which chimera filtering was conducted with--uchime_denovo. By focusing on centroids, this step reduced dataset size and computational demand, though at the expense of some sensitivity to rare chimeras; however, additional quality control measures, including abundance filtering and taxonomic validation, compensated for potential artifacts.

To reduce the impact of mitochondrial pseudogenes (NUMTs), several complementary filtering strategies were applied: de novo chimera detection and removal, length-based filtering, and taxonomic validation using BLAST searches against both a custom BOLD database and GenBank, combined with classification using the RDP classifier. Taxonomic assignments were refined through a consensus approach to minimize misclassification. Furthermore, we employed a negative-control-based filtering procedure: OTUs detected in biological samples with read counts not exceeding the maximum observed in negative controls were excluded. This conservative step mitigates spurious signals from NUMTs, cross-contamination, or environmental DNA artifacts. Given these layers of quality control, we are confident that NUMTs did not substantially inflate taxonomic diversity estimates.

Non-chimeric sequences were clustered into OTUs by means of SWARM v3.1.0 and the parameters-d 12-z, and an OTU table was generated by mapping reads back to OTUs using the VSEARCH--usearch_global utility. To minimize false positives, OTUs were verified against GenBank and BOLD databases using BLAST, following similarity and alignment quality criteria. Searches were performed locally with NCBI BLAST+ (blastn; ftp://ftp.ncbi.nlm.nih.gov/blast/executables/blast+/) using the following parameters:-task megablast-max_target_seqs 1-max_hsps 1-evalue 10-word_size 28-outfmt “6 sacc pident salltitles qseqid length evalue qcovs staxids”.

For each OTU, only the top hit by percentage identity was retained. Two databases were queried: (1) the NCBI nt database (downloaded from ftp://ftp.ncbi.nlm.nih.gov/blast/db/; download date reported in results), and (2) a custom-formatted BLAST database built from COI records obtained via a BOLD data package release (http://doi.org/10.5883/DP-BOLD_Public.27-Dec-2024). Most records lacking BINs were filtered out, except for rare taxa absent from any BIN-linked records. BIN assignments were made based on best hits to this BOLD database, ensuring control over query parameters and annotation consistency.

Taxonomic classification was further supported by the RDP Bayesian classifier (Wang et al. 2007), trained on a COI reference dataset (Porter and Hajibabaei 2018) to complement BLAST-based assignments. The RDP classifier provided bootstrap support values at each taxonomic rank, offering a probabilistic measure of assignment confidence. BLAST results from GenBank and BOLD were integrated with RDP outputs under a least common ancestor (LCA) consensus framework, ensuring conservative and robust classifications, particularly for degraded sequences or incomplete reference databases. Final taxonomic summaries were visualized using KronaTools v1.311.

## Statistical analysis

Temperature datasets were created using all temperature loggers, i.e., 72 Hobo vertical loggers and 152 horizontal Tomst loggers. The following temperature aspects were calculated using the 30-, and 15-minute logging intervals: daily average temperature, daily maximum temperature, daily daytime average temperature, and daily nighttime average temperature, and the standard deviation of all of these (heterogeneity). All daily temperature aspects were then summarized for each study plot. To assess temperature heterogeneity differences between forest management interventions (Gap and Thinning), generalized linear mixed models were used with the glmmTMB extension (Brooks 2017). Temperature heterogeneity was used as the response variable (of each temperature aspect) with management type as a predictor variable and study plot as a random effect. Temporal autocorrelation was accounted for with an *ou*() covariance structure to match the irregular timing of arthropod sampling events.

First, the arthropod dataset was further filtered using the “singleton filter” function (developed in Kortmann et al. 2025) in order to filter sequencing errors and real singletons (Chiu 2016). Then, arthropod samples were standardized for coverage, especially to account for the uneven sampling events due to either missing samples or the fact that some samples were exposed for slightly different time periods. Sample coverage standardization is a robust way to account for uneven samples (Gotelli and Colwell 2001, Roswell et al. 2021), where diversity and community metrics are either rarefied or extrapolated, depending on the selected coverage value. We used the package *iNEXT.beta3D* (Chao et al. 2023) to standardise our samples to a sample coverage = 0.9. After the standardization, community metrics was calculated for every sample-pair taking OTU abundance into account, along the Hill-numbers (Hill 1973, Chao et al. 2014). Using OTU read data; we can assess diversity from rare species to dominant species by adjusting the diversity order *q* when creating similarity matrices by taking relative abundances into account. For *q* = 0, the Hill number reduces to species richness and thus is more sensitive to rare species in the assemblage or sample; the measures with *q* =1 and *q* = 2 can be interpreted as the effective number of common and dominant species in the assemblage, respectively. This way, all rarity classifications are sample-dependent, as categories were assigned to species compared to others within the same assemblage. However, it is important to note that OTU read count cannot be directly translated to abundance data, therefore caution needs to be taken when investigating different aspects of diversity metrics based on OTU read counts. Still, we assume that for whole – community assessments, data based on read counts should represent the true patterns of response. Pairwise community similarity matrices were constructed using the *iNEXTbeta3D_pair3D* function from the *iNEXT*.*beta3D* package (Chao et al. 2023). For distance matrices in communities focusing on q = 0, Sorensen distance was used, for q = 1, Horn distance was used and for q = 2, Morisita-Horn distance was used. Arthropod communities were visualised along season, trap height and geographic location with a Nonmetric Multidimensional Scaling (NMDS) test using the *metaMDS* function from package *vegan* (Oksanen et al. 2015).

### Community similarity analysis

In order to assess how temperature similarity is driving arthropod community similarity, we created similarity matrices of all temperature aspects and additionally included a temperature index where daily averages, maximums, daytime averages, daytime maximums, nighttime averages and nighttime maximums were combined. To account for other environmental variables, we created similarity matrices for season (trap collection date), vertical position of arthropod trap (height from ground) and geographical location (based on the coordinates of single traps). All similarity matrices were created and standardised using the *decostand* function from the *vegan* package (Oksanen et al. 2015), using eucledian distances. Arthropod and environmental similarity matrices were then tested using Multiple Regression on Distance Matrices (MRM) from the *ecodist* package (Goslee and Urban 2007). All temperature aspects were tested separately due to correlation but visualised in one single graph for better clarity. Single temperature aspect similarity matrices were always combined with the relevant environmental matrices to account for all variables driving arthropod communities, however given that forest management type was driving temperature heterogeneity, Gap and Thinning was not included in the MRM analysis. Temperature similarity is assumed to highly represent vegetation and management type, therefore no environmental variable for these metrics were analysed.

## Results

Canopy temperature heterogeneity was higher at study plots where Gap treatment was applied than at study plots where Thinning treatment was applied, however the difference was only significant for temperature maximum heterogeneity and marginally significant for average temperature heterogeneity (Fig. 2). The presence of standing deadwood did not impact temperature heterogeneity (Supp. Table 1).

**Figure 2.**
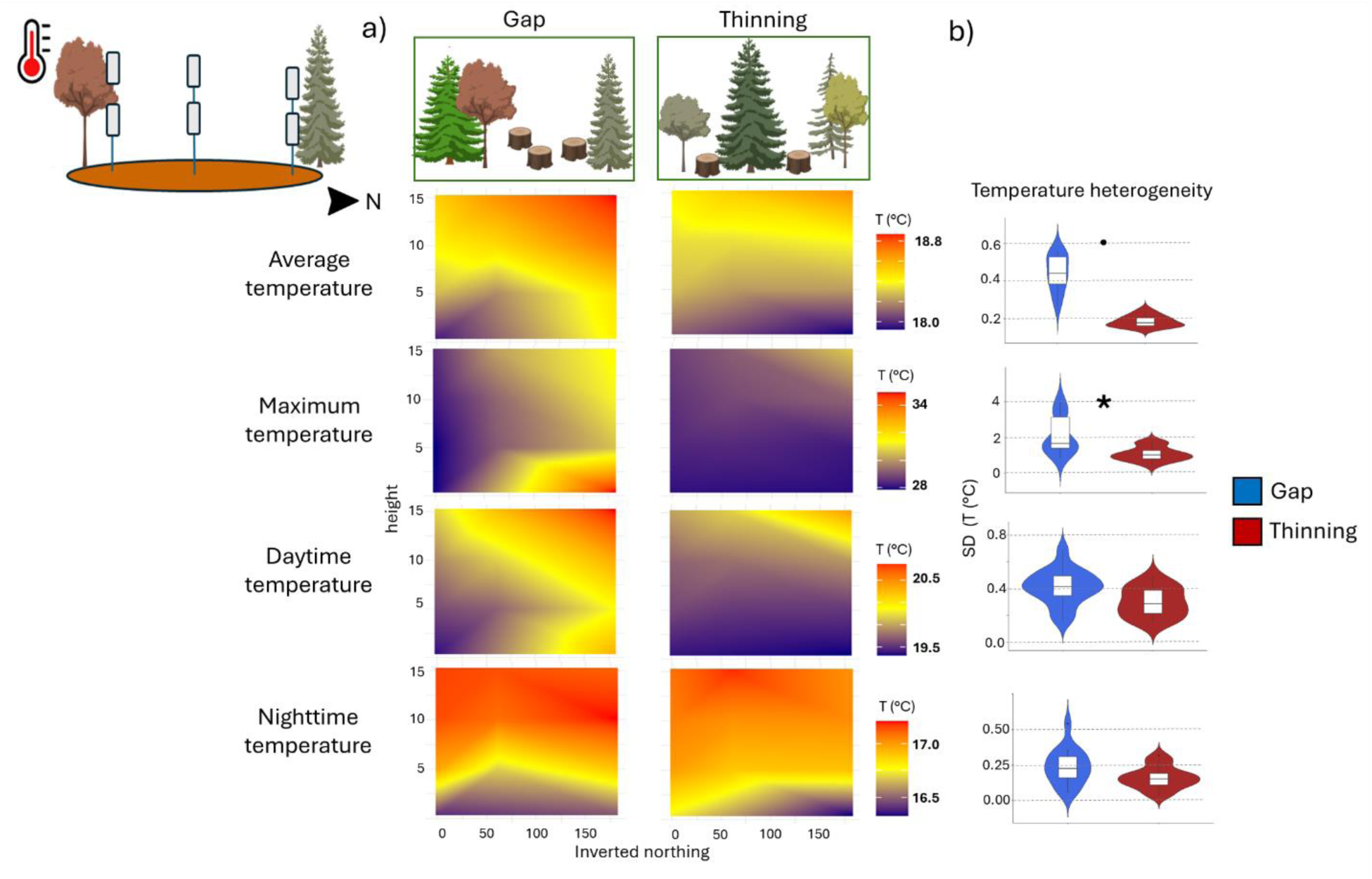
a) Projected *canopy* temperature heatmaps along the 4 height categories of the measured average, maximum, daytime and nighttime temperatures showing the fine-scale temperature ‘fingerprint’ of the two studied management types. Based on the fine-scale temperature measurements, b) temperature heterogeneity was calculated (standard deviation within each study plot) and shows differences between Gap (blue) and Thinning (red) treatments: point indicates marginally significant result (p < 0.1) and asterisk indicates significant impact (p < 0.05) of management type.

Forest floor temperature heterogeneity was significantly higher at study plots with Gap treatment for all temperature aspects (average, maximum, daytime, nighttime, Fig. 3). Standing deadwood did not have an impact on temperature heterogeneity (Supp. Table 1).

**Figure 3.**
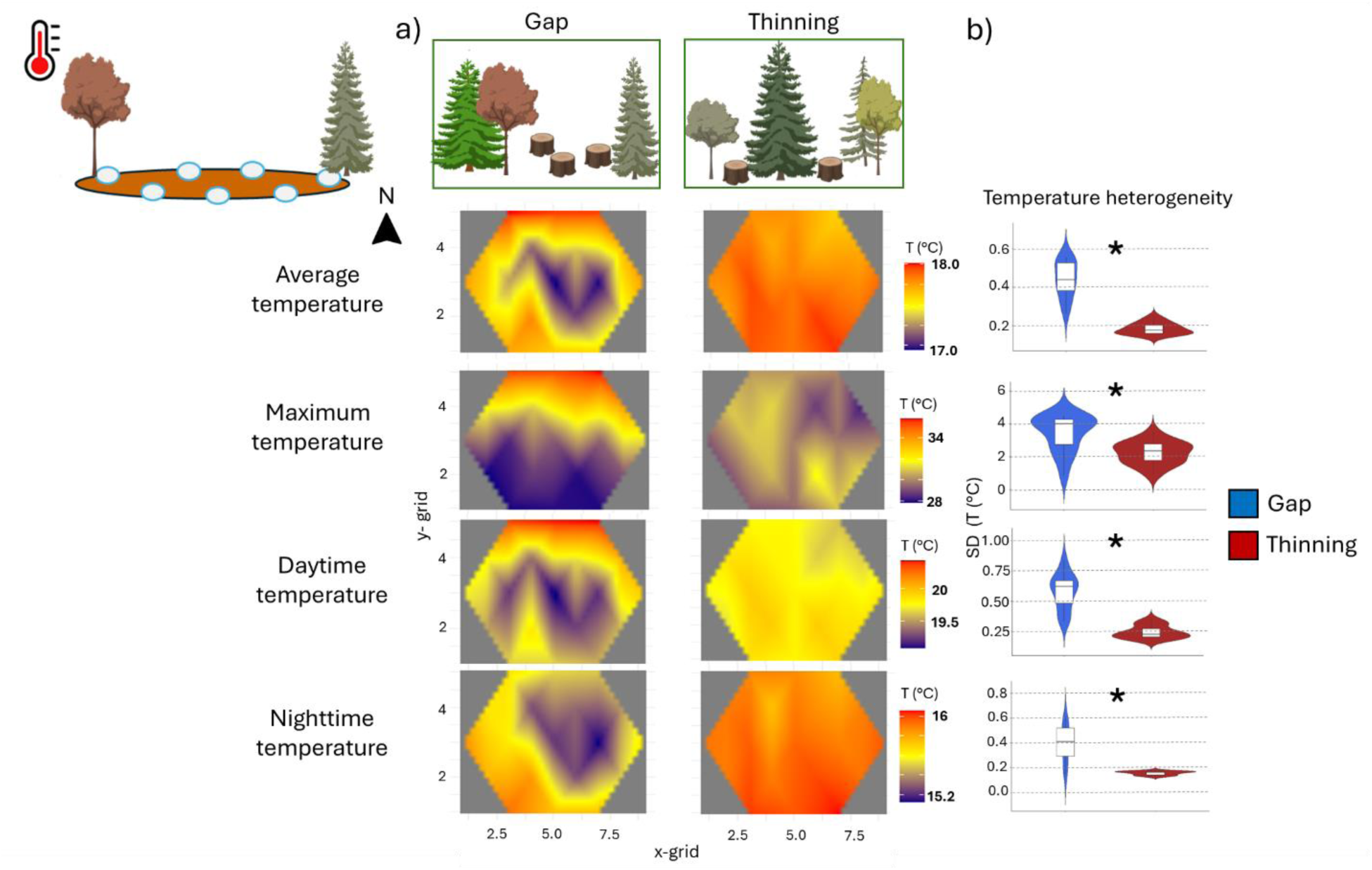
a) Projected *forest floor* temperature heatmaps along the 4 height categories of the measured average, maximum, daytime and nighttime temperatures showing the fine-scale temperature ‘fingerprint’ of the two studied management types. Based on the fine-scale temperature measurements, b) temperature heterogeneity was calculated (standard deviation within each study plot) and shows differences between Gap (blue) and Thinning (red) treatments: asterisk indicates significant impact (p < 0.05) of management type.

The metabarcoding of arthropod samples recovered 10,640 Operational Taxonomic Units (OTUs) in 44 Orders in the Phylum Arthropoda with 51% assigned to a genus and further 24% to species level, resulting in 2447 assigned arthropod species. After applying the singleton filtering step to remove sequencing errors, but retain rare OTUs in the dataset, 9,153 Arthropod OTUs remained in our final dataset. In flight-interception traps, Diptera, Hymenoptera and Coleoptera, and in pitfall traps, Coleoptera, Diptera and Entomobryomorpha orders gained the highest counts of OTU reads (Fig. 4).

**Figure 4.**
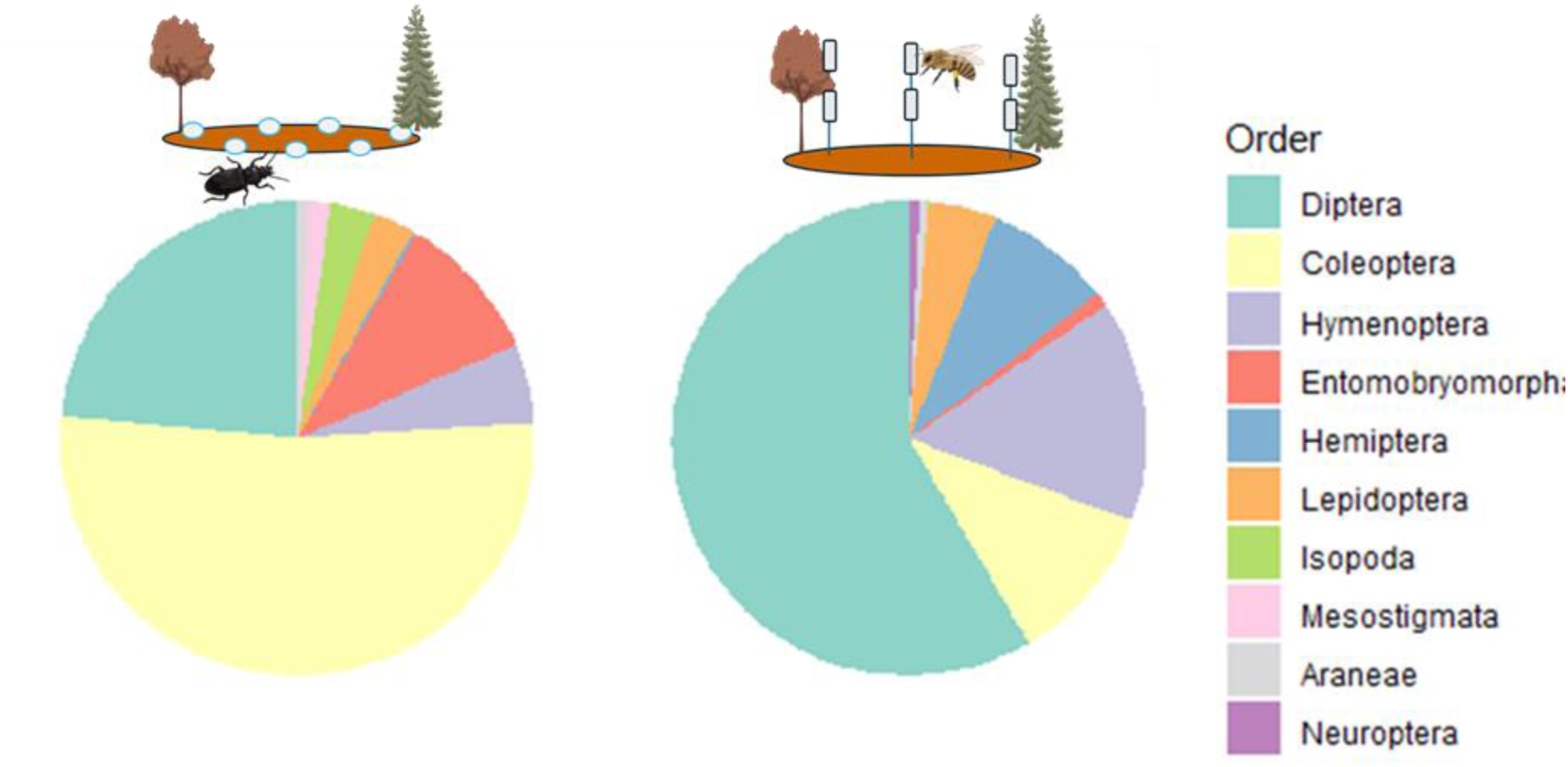
Proportions of read counts in pitfall (left) and flight-interception traps (right) of the ten taxonomic order which had the highest species read count.

The multiple regression on distance matrices (MRM) analysis revealed that overall arthropod community similarity is significantly affected by combined temperature metric and nighttime temperature similarity (Fig. 4a), but not all temperature similarity indices. The result was independent from rarity, i.e. *q*=0, *q*=1 and *q*=2 communities had the same pattern in response to temperature similarity. Trap height and season were both more important to drive arthropod community similarity than temperature similarity, and geographical location was also significant, but less important than temperature (Fig. 5a). However, given that the community similarity analysis revealed that arthropods communities separate based on trap-type (at height 0 m and other than 0 m, Supp. Fig. 1), we tested community similarity against temperature similarity for each trap types. This analysis revealed that arthropod communities inhabiting the canopy (recovered from flight-interception traps) focusing on rare species were affected by daily temperature maximum similarity. Season was the most important variable, while coefficients for height and location were similar to temperature. However, when focusing on communities of common and dominant species in the canopy, nighttime temperature similarity became a strong predictor of arthropod community similarity together with our combined daily temperature index. Season was always the most important variable, however with coefficients similar to nighttime temperature. Height similarity of the traps was not significantly impacting arthropod community similarity for common and dominant species (Fig. 5b).

**Figure 5.**
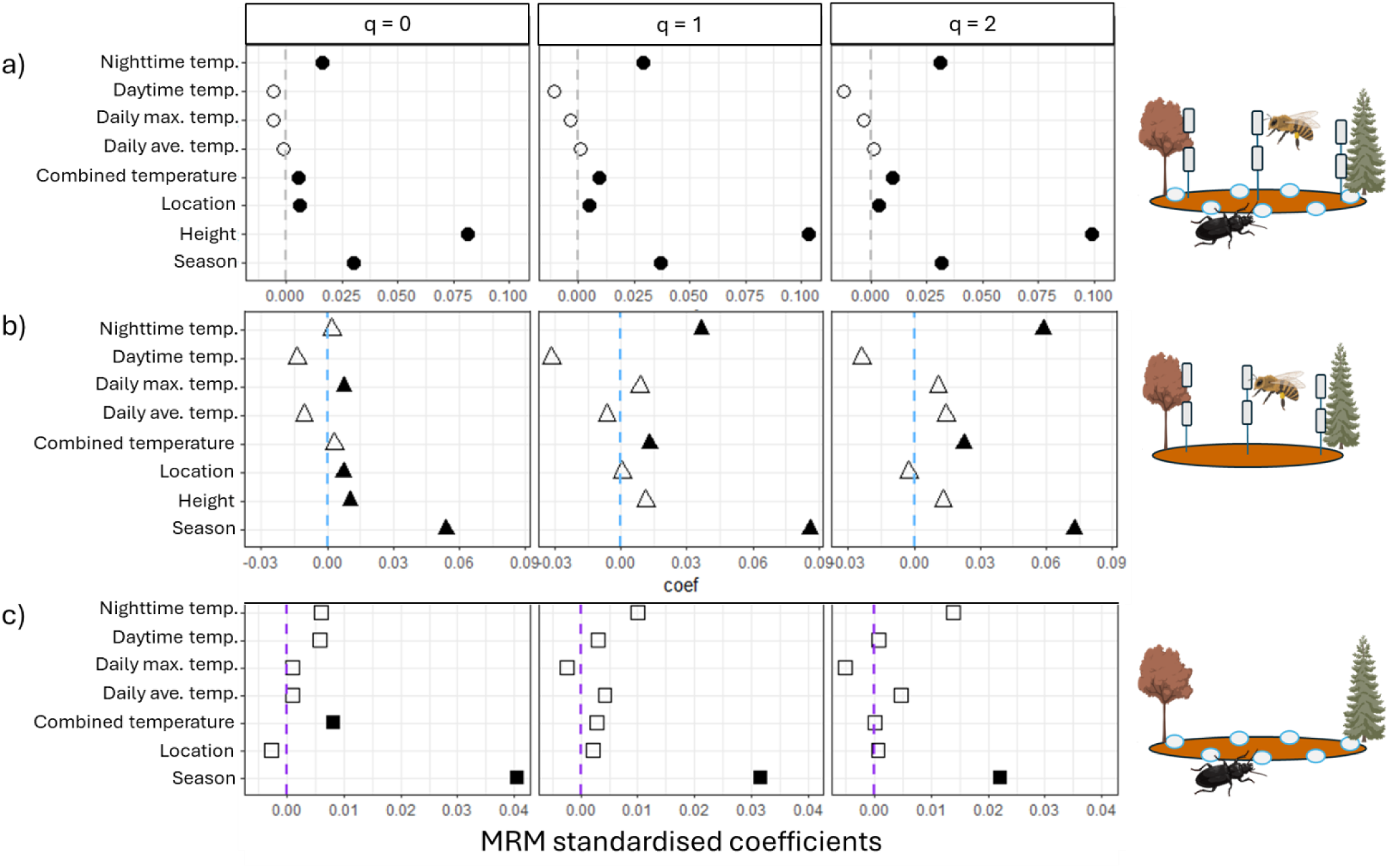
Output of MRM analysis, where arthropod community metrics were standardised by sample coverage and environmental metrics were standardised to calcualte eucledian distances. Panel a) shows *overall arthropod* community similarity responses to season, height, location and temperature metrics; while panel b) shows arthropod communities recovered from flight-interception traps from the vertical layer of the *canopy*; and panel c) shows arthropod communities recovered from pitfall traps, representing the horizontal layer of the *forest floor*. Filled symbols show a significant impact of a given variable on arthropod community similarity.

Arthropod communities recovered from pitfall traps were mostly responding to season (trapping date), regardless of rarity. Out of all temperature metrics, arthropod community similarity focusing on rare species were affected only by our combined daily temperature index (Fig. 5c).

Seasonality (similarity) was always the strongest predictor of arthropod community similarity, and temperature is also a variable that changes with seasons. We included both variables in our MRM analysis, because we believe that both predictors impact arthropod communities. This way some temperature variables could have been superseded by the season variable, as in some periods dates were changing with temperatures concurrently. However, ignoring sampling dates would have deleted an important factor in arthropod community seasonal changes.

## Discussion

There is a growing need in conservation to synthetise whole community responses to disturbance and management with microclimatic conditions in order to improve biodiversity conservation and restoration efforts (Sears et al. 2011, Sallé et al. 2021, Kemppinen et al. 2024). Our study shows that some aspect of fine-scale temperature drives overall arthropod community similarity in a temperate Central-European managed forest ecosystem.

Forests provide safe environments for organisms, via buffering climate extremes and other anthropogenic effects (Kulakowski et al. 2017), however native forest suffer from various mismanagement and climate-induced dieback (Allen et al. 2015, Hammond et al. 2022). It has been shown that canopy openings increase the temperature of a forest plot when the buffering capacity of the canopy is no longer present (Zellweger et al. 2019, Horváth et al. 2023). This could result in “winner or loser” species and shape communities, depending on the severity and area of the canopy loss (Sire et al. 2022). Even though large canopy gaps could be detrimental for ground-dwelling arthropods via increasing ground temperatures (Hartshorn 2021, Barahona-Segovia et al. 2022), when established on relatively small areas, the created microclimate heterogeneity increases arthropod (and other) diversity (Kriegel et al. 2021, Lettenmaier et al. 2022, Rothacher et al. 2025). However, describing fine-scale temperature dynamics within forest plots are only recently gaining attention to show the dramatic impact of canopy manipulation on the microclimates of different forest layers (Frey et al. 2023, Horváth et al. 2023). Our study shows that beyond diversity (Pierick et al., 2025), whole communities also respond to microclimate and temperature similarity.

It is important to not only measure average temperatures, but components of daily temperatures, as they could shape microclimates differently and changes might occur asymmetrically throughout the daily cycle, with increased fluctuations affecting organisms more profoundly (Terblanche et al. 2010, Paaijmans et al. 2013). Moreover, time of activity differs between arthropod taxa; therefore, communities could respond differently to temperatures based on their daily activity patterns. Interestingly, based on historic datasets, overall average temperatures are increasing via nighttime temperatures at a faster rate than daytime temperatures (Vose et al. 2005, Birch 2014, Davy et al. 2017). Furthermore, studies investigating responses to temperature changes are most often focusing on physiological responses or performance of single species to warming (Ma et al. 2004, McMillan et al. 2005, Estay et al. 2011, Piyaphongkul et al. 2012, Ribeiro et al. 2012). Here, our study is the first to show that besides combined daily temperature index, nighttime microclimatic differences could drive whole forest arthropod communities, regardless of their rarity within the community. Low nighttime temperatures are regarded as a recovery period following a period when heat stress might have occurred during the day (Colinet et al. 2015), therefore with increased nighttime temperatures, this is limited, and reduces the nymphal survival and adult performance in aphids (Zhao et al. 2014). Our results imply that nighttime temperature might not only affect the physiology of individual species, but also play a role in shaping whole arthropod communities in temperate forests. Species might disappear or appear in local communities based on how nighttime temperatures changes following days with increased daytime temperatures. When nighttime is no longer a recovery period for some arthropods, only warm-associated species might be able to maintain their populations in localities, where temperatures do not decrease to an optimal temperature during the night. This might have been also reflected in the significant response of overall community similarity to the combined weather variable, where both daytime and nighttime averages and maximums were included. This variable could be a proxy to daily temperature variation; showing that species might be filtered in communities according to daytime and nighttime temperature fluctuations.

Arthropods inhabiting the canopy of a forest are a highly diverse group which are often specialised in specific behaviours (foraging strategies, host-tree relationships, breeding patterns, etc. (Nakamura et al. 2017, Ruchin 2023)). They are also the most sensitive group to forest management strategies as forestry activities most often remove whole trees and heavily alter canopy structure (Stork et al. 1997). Stratification is documented in tropical and temperate forests (Basset 2003, Nakamura et al. 2017, de Souza Amorim et al. 2022), but was rarely linked to temperature differences, even though canopy opening affects temperatures significantly. Arthropod communities inhabiting the canopy showed a strong response to nighttime temperature similarity, even more than to height, but only communities weighed for common and dominant species, while communities weighed for rare species were not responding to nighttime temperature similarity, but daily maximum temperature similarity. Canopy openness is a strong driver of forest arthropods (Thorn et al. 2016, Černecká et al. 2020, Cours et al. 2023), and this variable is dramatically changing maximum temperatures (Brůna et al. 2024). Therefore, forest-affiliated species which are likely to be cold-associated, temperature maximum similarities are likely to shape community similarity within these rare forest assemblages.

Ground-dwelling arthropod community similarities were largely not affected by temperature similarities. Only communities weighed for rare species were driven by our combined daily temperature variable. This might indicate that there is an extremely high variation of available ground structures, and ground temperatures which our study might have failed to detect. We measured ground temperatures and arthropod communities with only seven pitfall traps and associated temperature loggers, which could have failed in describing the full heterogeneity of the ground habitats and microclimates. Together with this, it could be possible that arthropods are responding to microhabitat heterogeneity attributes together (such as resources, enemy-free or competition-free spaces, species interactions, etc.), not only the temperature of the microhabitat. Perhaps ground-dwelling arthropods are foraging in all microhabitats depending on the time of the day and the season, that our temporal resolution was too low to detect temperature preferences of a whole community. Season (date of sample collection) was always the strongest predictor of both canopy and ground inhabiting arthropod communities, therefore sampling more often could overcome the significant seasonal impacts. Future studies could focus more not only on spatially fine-scale temperature and community metrics, but also temporally fine-scale data collection. This way it should be possible to highlight the exact responses of temperature fluctuations and responses to any extreme events.

## Conclusions

Given the many recent advantages in modelling, and temperature logging methods (summarised in Kemppinen et al. 2024), and the advancement of DNA-based species identification, monitoring both microclimates and whole arthropod communities is possible. When establishing links between management (or disturbance), microclimates and species communities, it is possible to create a way forward in management for species conservation. Our dataset sheds a light on whole-arthropod community responses to different aspects of temperature, but more studies are needed to understand the process and predict future community shifts. The detected patterns might change depending on the resolution used in our approach, and these fine-scale differences might be more pronounced on a lower resolution or might diminish. The canopy and floor of a forest are both constantly being modified via management interventions, and monitoring temperature dynamics which alter arthropod species and communities should be integrated in management decisions and help develop optimal interventions.

