## Supplementary Information for "Forest vertical and horizontal temperature similarity drives arthropod communities in a managed temperate forest"

Supplementary results

**Supplementary Table 1**. The table shows the outcome of generalised linear models with an extension to correct for temporal autocorrelation (glmmTMB model family). Bold characters indicate a significant impact.

| Logger type | Temperature | Variable | Chi-squared | p - value |
| --- | --- | --- | --- | --- |
| 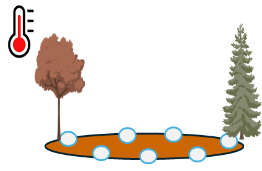 | Daily mean | **Mngmnt** | **55.64** | **> 0.001** |
|  |  | Standing deadwood | 0.41 | 0.520 |
|  |  | Interaction | 0.76 | 0.382 |
|  | Daily max. | **Mngmnt** | **4.46** | **0.034** |
|  |  | Standing deadwood | 0.25 | 0.617 |
|  |  | Interaction | 0.01 | 0.921 |
|  | Daytime average | **Mngmnt** | **25.20** | **> 0.001** |
|  |  | Standing deadwood | 0.01 | 0.940 |
|  |  | Interaction | 0.13 | 0.708 |
|  | Nighttime average | **Mngmnt** | **12.82** | **> 0.001** |
|  |  | Standing deadwood | 0.93 | 0.333 |
|  |  | Interaction | 0.44 | 0.505 |
| 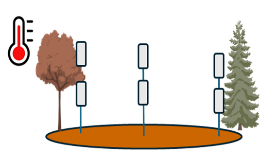 | Daily mean | *Mngmnt* | *3.61* | *0.057* |
|  |  | Standing deadwood | 0.05 | 0.816 |
|  |  | Interaction | 1.45 | 0.227 |
|  | Daily max. | **Mngmnt** | **4.49** | **0.033** |
|  |  | Standing deadwood | 4.48 | 0.484 |
|  |  | Interaction | 0.22 | 0.637 |
|  | Daytime average | Mngmnt | 3.30 | 0.069 |
|  |  | Standing deadwood | 0.21 | 0.648 |
|  |  | Interaction | 0.64 | 0.422 |
|  | Nighttime average | Mngmnt | 1.83 | 0.176 |
|  |  | Standing deadwood | 0.01 | 0.917 |
|  |  | Interaction | 0.83 | 0.657 |

**Supplementary Table 2.** The table shows the outcome of each MRM model testing environmental and temperature variables using the whole arthropod community. The outcome includes standardised coefficient and significance test along the Hill-numbers (q = 0, q = 1, q = 2). Negative coefficients with high significance were ignored, as they make no ecological sense and are likely to be mathematical artifacts.

| 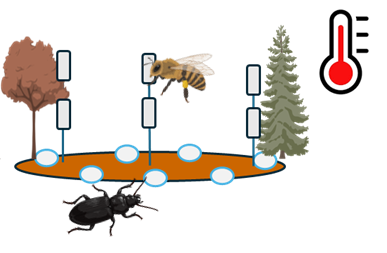 | Variable | Standard coefficient | p-value |
| --- | --- | --- | --- |
|  | **q = 0** | | |
|  | Intercept | 0.682 | 0.001 |
|  | Season | 0.030 | 0.001 |
|  | Height | 0.081 | 0.001 |
|  | Location | 0.006 | 0.001 |
|  | Combined daily temperature | 0.005 | 0.010 |
|  | Intercept | 0.693 | 0.001 |
|  | Season | 0.033 | 0.001 |
|  | Height | 0.083 | 0.00 |
|  | Location | 0.006 | 0.001 |
|  | Average daily temperature | -0.001 | 0.687 |
|  | Intercept | 0.699 | 0.001 |
|  | Season | 0.032 | 0.001 |
|  | Height | 0.083 | 0.001 |
|  | Location | 0.006 | 0.001 |
|  | Daily maximum temperature | -0.005 | 0.037 |
|  | Intercept | 0.696 | 0.001 |
|  | Season | 0.034 | 0.001 |
|  | Height | 0.083 | 0.001 |
|  | Location | 0.006 | 0.001 |
|  | Daytime average temperature | -0.005 | 0.018 |
|  | Intercept | 0.683 | 0.001 |
|  | Season | 0.028 | 0.001 |
|  | Height | 0.078 | 0.001 |
|  | Location | 0.006 | 0.001 |
|  | Nighttime average temperature | 0.016 | 0.001 |
|  | **q = 1** | | |
|  | Intercept | 0.644 | 0.999 |
|  | Season | 0.037 | 0.001 |
|  | Height | 0.103 | 0.001 |
|  | Location | 0.005 | 0.001 |
|  | Combined daily temperature | 0.009 | 0.002 |
|  | Intercept | 0.661 | 0.999 |
|  | Season | 0.041 | 0.001 |
|  | Height | 0.107 | 0.001 |
|  | Location | 0.005 | 0.001 |
|  | Average daily temperature | 0.001 | 0.742 |
|  | Intercept | 0.665 | 0.999 |
|  | Season | 0.041 | 0.001 |
|  | Height | 0106 | 0.001 |
|  | Location | 0.005 | 0.001 |
|  | Daily maximum temperature | -0.003 | 0.326 |
|  | Intercept | 0.669 | 0.001 |
|  | Season | 0.044 | 0.001 |
|  | Height | 0.107 | 0.001 |
|  | Location | 0.005 | 0.001 |
|  | Daytime average temperature | -0.010 | 0.003 |
|  | Intercept | 0.645 | 0.001 |
|  | Season | 0.034 | 0.001 |
|  | Height | 0.098 | 0.001 |
|  | Location | 0.005 | 0.001 |
|  | Nighttime average temperature | 0.028 | 0.001 |
|  | **q = 2** | | |
|  | Intercept | 0.673 | 0.999 |
|  | Season | 0.032 | 0.001 |
|  | Height | 0.099 | 0.001 |
|  | Location | 0.003 | 0.001 |
|  | Combined daily temperature | 0.009 | 0.002 |
|  | Intercept | 0.691 | 0.999 |
|  | Season | 0.036 | 0.001 |
|  | Height | 0.102 | 0.001 |
|  | Location | 0.004 | 0.001 |
|  | Average daily temperature | 0.001 | 0.796 |
|  | Intercept | 0.695 | 0.999 |
|  | Season | 0.036 | 0.001 |
|  | Height | 0.102 | 0.001 |
|  | Location | 0.004 | 0.001 |
|  | Daily maximum temperature | -0.003 | 0.461 |
|  | Intercept | 0.700 | 0.999 |
|  | Season | 0.040 | 0.001 |
|  | Height | 0.103 | 0.001 |
|  | Location | 0.004 | 0.001 |
|  | Daytime average temperature | -0.012 | 0.004 |
|  | Intercept | 0.674 | 0.999 |
|  | Season | 0.028 | 0.001 |
|  | Height | 0.094 | 0.001 |
|  | Location | 0.004 | 0.001 |
|  | Nighttime average temperature | 0.031 | 0.001 |

**Supplementary Table 3.** The table shows the outcome of each MRM model testing environmental and temperature variables using the only the canopy arthropod community (flight-interception traps). The outcome includes standardised coefficient and significance test along the Hill-numbers (q = 0, q = 1, q = 2). Negative coefficients with high significance were ignored, as they make no ecological sense and are likely to be mathematical artifacts.

| 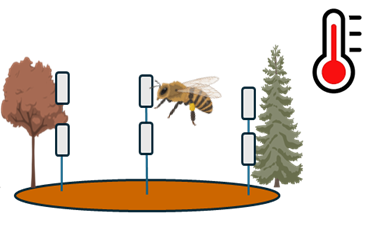 | Variable | Standard coefficient | p-value |
| --- | --- | --- | --- |
|  | **q = 0** | | |
|  | Intercept | 0.722 | 0.682 |
|  | Season | 0.054 | 0.001 |
|  | Height | 0.010 | 0.001 |
|  | Location | 0.007 | 0.001 |
|  | Combined daily temperature | 0.003 | 0.166 |
|  | Intercept | 0.733 | 0.028 |
|  | Season | 0.062 | 0.001 |
|  | Height | 0.010 | 0.001 |
|  | Location | 0.007 | 0.001 |
|  | Average daily temperature | -0.010 | 0.003 |
|  | Intercept | 0.720 | 0.794 |
|  | Season | 0.056 | 0.001 |
|  | Height | 0.010 | 0.001 |
|  | Location | 0.007 | 0.001 |
|  | Daily maximum temperature | 0.007 | 0.027 |
|  | Intercept | 0.720 | 0.002 |
|  | Season | 0.062 | 0.001 |
|  | Height | 0.011 | 0.001 |
|  | Location | 0.007 | 0.001 |
|  | Daytime average temperature | -0.014 | 0.001 |
|  | Intercept | 0.727 | 0.343 |
|  | Season | 0.055 | 0.001 |
|  | Height | 0.010 | 0.001 |
|  | Location | 0.007 | 0.001 |
|  | Nighttime average temperature | 0.002 | 0.558 |
|  | **q = 1** | | |
|  | Intercept | 0.646 | 0.999 |
|  | Season | 0.086 | 0.001 |
|  | Height | 0.011 | 0.081 |
|  | Location | 0.001 | 0.414 |
|  | Combined daily temperature | 0.012 | 0.045 |
|  | Intercept | 0.674 | 0.999 |
|  | Season | 0.099 | 0.001 |
|  | Height | 0.012 | 0.072 |
|  | Location | 0.001 | 0.232 |
|  | Average daily temperature | -0.006 | 0.501 |
|  | Intercept | 0.661 | 0.999 |
|  | Season | 0.095 | 0.001 |
|  | Height | 0.012 | 0.087 |
|  | Location | 0.001 | 0.306 |
|  | Daily maximum temperature | 0.008 | 0.327 |
|  | Intercept | 0.688 | 0.999 |
|  | Season | 0.110 | 0.001 |
|  | Height | 0.013 | 0.043 |
|  | Location | 0.001 | 0.081 |
|  | Daytime average temperature | -0.031 | 0.001 |
|  | Intercept | 0.655 | 0.999 |
|  | Season | 0.073 | 0.001 |
|  | Height | 0.011 | 0.068 |
|  | Location | 0.001 | 0.262 |
|  | Nighttime average temperature | 0.036 | 0.001 |
|  | **q = 2** | | |
|  | Intercept | 0.649 | 0.999 |
|  | Season | 0.072 | 0.001 |
|  | Height | 0.013 | 0.106 |
|  | Location | -0.002 | 0.046 |
|  | Combined daily temperature | 0.022 | 0.004 |
|  | Intercept | 0.687 | 0.999 |
|  | Season | 0.079 | 0.001 |
|  | Height | 0.014 | 0.082 |
|  | Location | -0.002 | 0.054 |
|  | Average daily temperature | 0.014 | 0.163 |
|  | Intercept | 0.685 | 0.999 |
|  | Season | 0.085 | 0.001 |
|  | Height | 0.014 | 0.074 |
|  | Location | -0.002 | 0.044 |
|  | Daily maximum temperature | 0.011 | 0.303 |
|  | Intercept | 0.713 | 0.998 |
|  | Season | 0.094 | 0.001 |
|  | Height | 0.015 | 0.049 |
|  | Location | -0.002 | 0.057 |
|  | Daytime average temperature | -0.023 | 0.023 |
|  | Intercept | 0.657 | 0.999 |
|  | Season | 0.061 | 0.001 |
|  | Height | 0.014 | 0.079 |
|  | Location | -0.001 | 0.126 |
|  | Nighttime average temperature | 0.058 | 0.001 |

**Supplementary Table 4.** The table shows the outcome of each MRM model testing environmental and temperature variables using the only the forest floor arthropod community (pitfall traps). The outcome includes standardised coefficient and significance test along the Hill-numbers (q = 0, q = 1, q = 2). Negative coefficients with high significance were ignored, as they make no ecological sense and are likely to be mathematical artifacts.

| 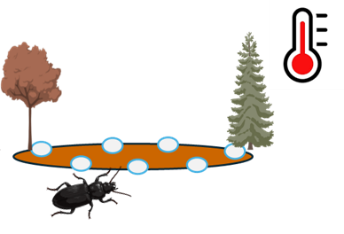 | Variable | Standard coefficient | p-value |
| --- | --- | --- | --- |
|  | **q = 0** | | |
|  | Intercept | 0.599 | 0.999 |
|  | Season | 0.040 | 0.001 |
|  | Location | 0.009 | 0.001 |
|  | Combined daily temperature | 0.008 | 0.033 |
|  | Intercept | 0.617 | 0.999 |
|  | Season | 0.043 | 0.001 |
|  | Location | 0.009 | 0.001 |
|  | Average daily temperature | 0.001 | 0.894 |
|  | Intercept | 0.617 | 0.999 |
|  | Season | 0.043 | 0.001 |
|  | Location | 0.009 | 0.001 |
|  | Daily maximum temperature | 0.001 | 0.306 |
|  | Intercept | 0.613 | 0.999 |
|  | Season | 0.042 | 0.001 |
|  | Location | 0.009 | 0.001 |
|  | Daytime average temperature | 0.005 | 0.306 |
|  | Intercept | 0.613 | 0.999 |
|  | Season | 0.042 | 0.001 |
|  | Location | 0.009 | 0.001 |
|  | Nighttime average temperature | 0.006 | 0.264 |
|  | **q = 1** | | |
|  | Intercept | 0.575 | 0.999 |
|  | Season | 0.031 | 0.001 |
|  | Location | 0.015 | 0.001 |
|  | Combined daily temperature | 0.002 | 0.585 |
|  | Intercept | 0.578 | 0.999 |
|  | Season | 0.031 | 0.001 |
|  | Location | 0.015 | 0.001 |
|  | Average daily temperature | 0.004 | 0530 |
|  | Intercept | 0.584 | 0.999 |
|  | Season | 0.032 | 0.001 |
|  | Location | 0.015 | 0.001 |
|  | Daily maximum temperature | -0.002 | 0.765 |
|  | Intercept | 0.579 | 0.999 |
|  | Season | 0.032 | 0.001 |
|  | Location | 0.015 | 0.001 |
|  | Daytime average temperature | 0.003 | 0.650 |
|  | Intercept | 0.573 | 0.999 |
|  | Season | 0.030 | 0.001 |
|  | Location | 0.015 | 0.001 |
|  | Nighttime average temperature | 0.009 | 0.132 |
|  | **q = 2** | | |
|  | Intercept | 0.629 | 0.954 |
|  | Season | 0.022 | 0.003 |
|  | Location | 0.011 | 0.001 |
|  | Combined daily temperature | 0.001 | 0.995 |
|  | Intercept | 0.626 | 0.998 |
|  | Season | 0.021 | 0.003 |
|  | Location | 0.011 | 0.001 |
|  | Average daily temperature | 0.005 | 0.653 |
|  | Intercept | 0.635 | 0.981 |
|  | Season | 0.022 | 0.004 |
|  | Location | 0.012 | 0.001 |
|  | Daily maximum temperature | -0.005 | 0.654 |
|  | Intercept | 0.629 | 0.991 |
|  | Season | 0.022 | 0.004 |
|  | Location | 0.011 | 0.001 |
|  | Daytime average temperature | 0.001 | 0.965 |
|  | Intercept | 0.617 | 0.999 |
|  | Season | 0.019 | 0.022 |
|  | Location | 0.012 | 0.001 |
|  | Nighttime average temperature | 0.014 | 0.183 |

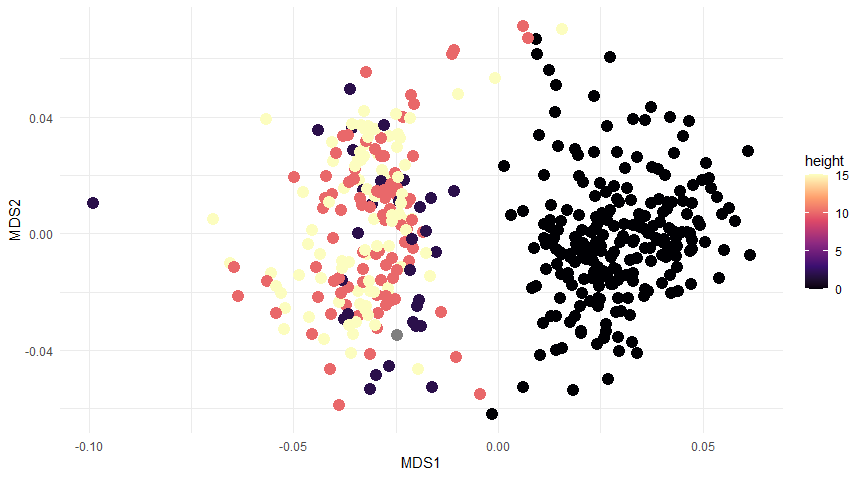

**Supplementary Figure 1**. NMDS ordination of arthropod communities along the height of the samples.

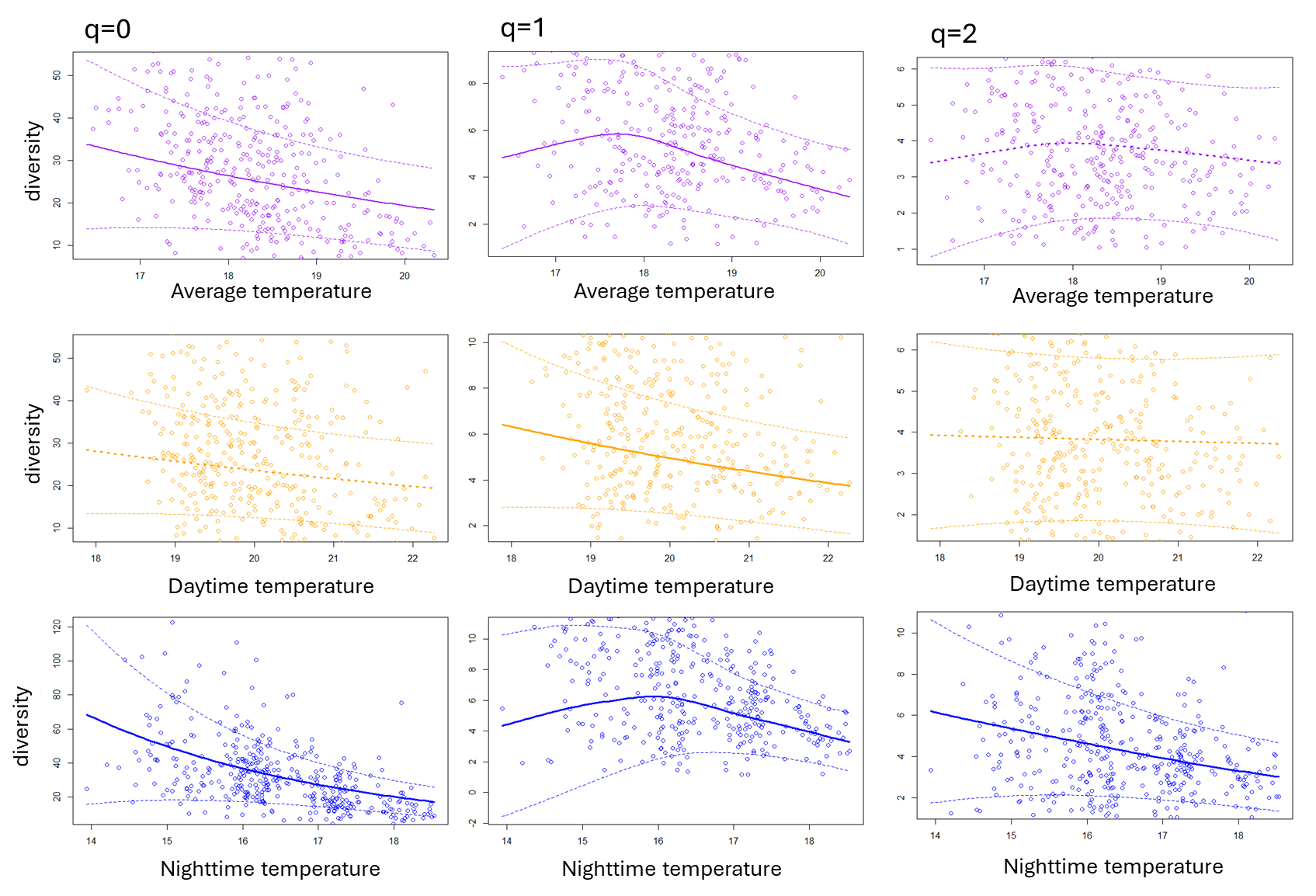

**Supplementary Figure 2**. Negative binomial model output of alpha diversity of arthropods driven by average, daytime and nighttime temperatures. Relationships with solid lines indicate a significant impact of temperature metric on alpha diversity.
